# Temporal Changes in Transcripts of MITE Transposable Elements during Rice Endosperm Development

**DOI:** 10.1101/2021.09.27.461986

**Authors:** Hiroki Nagata, Akemi Ono, Kaoru Tonosaki, Taiji Kawakatsu, Kentaro Yano, Yuji Kishima, Tetsu Kinoshita

**Author notes:** **Author contributions:** H.N.; A.O.; K.T.; T.Ki. conceived the research; A.O.; K.T.; T.Ka.; K.Y. supervised the experiments; H.N. performed all experiments; Y.K. provided materials.; H. N. drafted the manuscript, designed the experiments and analyzed the data; A.O.; T.Ki. helped design experiments and revise the draft; T.Ki. wrote the article with contributions from all authors.

## Abstract

The repression of transcription from transposable elements (TEs) by DNA methylation is necessary to maintain genome integrity and prevent harmful mutations. However, under certain circumstances, TEs are thought to escape from the host defense system and reactivate their transcription. In Arabidopsis (*Arabidopsis thaliana*) and rice (*Oryza sativa*), DNA demethylases target the sequences derived from TEs in the central cell, the progenitor cell for the endosperm in the female gametophyte. This genome-wide DNA demethylation is also observed in the endosperm after fertilization. In this study, we used a custom microarray to survey the transcripts generated from TEs during the rice endosperm development and at selected timepoints in the embryo as a control. The expression patterns of TE transcripts are dynamically up- and downregulated during endosperm development, especially for miniature inverted-repeat transposable elements (MITEs). Surprisingly, some TE transcripts were directionally controlled, while the other DNA transposons and retrotransposons were not. We also discovered the NF-Y binding motif, CCAAT, in the region near the 5′ terminal inverted repeat of *Youren*, one of the transcribed MITEs in the endosperm. Our results uncover dynamic changes in TE activity during endosperm development in rice.

## INTRODUCTION

The transcription of autonomous transposable elements (TEs) is essential for the transposition of class I RNA retrotransposons and class II DNA transposons (Wells and Feschotte, 2020) but is typically silenced by epigenetic mechanisms such as DNA methylation (Zhang et al., 2018). DNA methylation also disarms non-autonomous elements via transcriptional repression of their RNA intermediate or by blocking transposase binding to its target sequence required for “cut-and-paste” activity (Gierl et al., 1988; Hashida et al., 2006). Although DNA methylation plays essential roles in the repression of TE activity, this epigenetic mark is passively removed from TE sequences during DNA replication or actively erased by DNA glycosylases during the plant life cycle and in various tissues, including seed endosperm (Bartels et al., 2018; Ono and Kinoshita, 2021).

The endosperm is the nutritive tissue that sustains embryonic growth during seed development and seedling growth after germination, especially in crop species such as rice (*Oryza sativa*) (Lopes and Larkins, 1993; Sabelli and Larkins, 2009). The endosperm arises from the second fertilization event between the central cell of the female gametophyte and one of the two sperm cells delivered by the male pollen tube (Berger et al., 2008). The embryo is itself the product of the fertilization of the egg and the other sperm cell. Unlike the embryo, the endosperm has unique characteristics; its genome comprises two maternal copies and one paternal copy, undergoes terminal differentiation, and is subjected to global DNA demethylation (Lauria et al., 2004; Hsieh et al., 2009; Zemach et al., 2010).

In Arabidopsis (*Arabidopsis thaliana*), one of the DNA glycosylase termed DEMETER (DME) targets AT-rich transposons in euchromatin (Gehring et al., 2006; Ibarra et al., 2012) as well as the heterochromatic TEs with the histone chaperone FACT in the central cell, the progenitor cell of the endosperm before fertilization (Ikeda et al., 2011; Frost et al., 2018). This active demethylation also contributes to parent-of-origin specific gene expression, known as imprinting, in the endosperm (Fujimoto et al., 2011; Hsieh et al., 2011), possibly via an association with TEs (Qiu and Kohler, 2020; Rodrigues et al., 2021). Indeed, many TEs are thought to be activated in the endosperm, which is consistent with pioneering work exploring the causes of kernel pigmentation patterns (Doring and Starlinger, 1986) and maternal DNA hypomethylation in maize (*Zea mays*) endosperm (Lauria et al., 2004). While the activity of TEs is typically repressed by their host defense system to prevent harmful mutations and maintain genome integrity, the global DNA demethylation observed in the endosperm constitutes a notable exception during plant development. The current explanation predicts that TE transcripts from the central cell or the endosperm are processed into small RNAs that are then transported to the egg cell and the embryo to reinforce the repression of TEs in their genomes for the next generation (Bourc’his and Voinnet, 2010). In addition, the endosperm is a terminally differentiated tissue, which might therefore tolerate high TE activity brought upon by DNA demethylation. De-repression of TE activity influences the neighboring genomic regions (Yang et al., 2005; Naito et al., 2009), making the endosperm an ideal tissue to dissect the roles of TEs in relation to transcriptional control and genome integrity.

The rice genome is relatively small (389 Mb) compared to those of other crop species, and only about three times larger than that of Arabidopsis (125 Mb) (Arabidopsis Genome Initiative, 2000; International Rice Genome Sequencing, 2005). However, rice contains much more repetitive sequences than does Arabidopsis (Song and Cao, 2017). The TE distribution is also different: TEs in Arabidopsis are mainly located in the pericentromeric regions, while those in rice also populate in the euchromatin (Mirouze and Vitte, 2014). A striking feature of the rice genome with regard to copy number and TE distribution is its enrichment in DNA transposons, especially miniature inverted-repeat transposable elements (MITEs) (Jiang et al., 2004). MITEs are internal deletion derivatives of autonomous DNA transposons with terminal inverted repeats (TIRs) at both ends (Feschotte et al., 2002; Feschotte and Wessler, 2002; Feng, 2003). These short stretches of sequences, 100–600 bp in length, mostly reside in the euchromatic regions and are relatively close to genes compared to longer long terminal repeat (LTR)–type retrotransposons (Oki et al., 2008; Zemach et al., 2010).

In this study, we performed a time-course transcriptome analysis of rice TEs during endosperm development. We employed a custom-made microarray designed to detect transcripts generated from repetitive sequences in rice (Ishiguro et al., 2014), to bypass the limitations associated with short-read sequencing of TEs, especially for short and high-copy number elements like MITEs. We determined that many expressed TEs in the endosperm tended not to be expressed in the embryo and showed a temporal dependent expression pattern. Among expressed TEs, many MITEs displayed a directional behavior, as we detected transcripts originating from one strand only, likely reflecting the direction of the ancestral autonomous transcriptional unit. This pattern was not present in other tested DNA transposons and retrotransposons. We also identified the binding motif for NUCLEAR FACTOR Y (NF-Y) transcription factors (Mantovani, 1999; Nardini et al., 2013) within *Youren* sequences, which overlapped with the boundary of the 5′ TIR and the internal region. Together, our results suggest that while MITEs are inserted proximal to genes or coding regions in random orientations, MITE transcripts do not hitchhike the directional transcription of their neighboring genes (i.e., readthrough or chimeric transcripts) but rather exhibit their own directional transcriptional control, possibly starting from intrinsic sequences in *Youren* MITEs. Such an active transcriptional environment at MITE sequences may contribute to shaping the transcriptional network of genes expressed in the rice endosperm.

## RESULTS

### Expression of TEs in the embryo and the endosperm

Genome-wide hypomethylation and TE transposition are prominent features of the endosperm, but how TE expression changes over time during seed development is not well described. Therefore, we conducted a transcriptome analysis with a custom 44k Agilent microarray (Ishiguro et al., 2014) harboring 31,366 repeat-related, 864 microRNA (miRNA)–related probes, as well as probes for 9,563 genes. To investigate the dynamics of TE transcription, we isolated total RNAs from rice endosperm in 24-h intervals from 2 to 7 days after pollination (DAP) and at 10 DAP (Supplemental Figure S1A). The endosperm became coenocytic at 2 DAP and grew in a centripetal direction by 3 DAP. Cellularization took place between these time points in our growth conditions (Tonosaki et al., 2021). The endosperm continued to divide and began to deposit storage compounds until the maturation stage at 10 DAP (Supplemental Figure S1A). Our chosen time points thus covered most developmental transitions in endosperm development. In addition to the endosperm, we also extracted total RNAs from developing embryos collected at 5, 7 and 10 DAP as controls, which correspond to the coleoptile, the completion of organogenesis, and the maturation stages of rice embryogenesis, respectively (Itoh et al., 2016) (Supplemental Figure S1B).

We performed a principal component analysis (PCA) using the expression of genes and TEs obtained from all samples across replicates. We observed separation of genes and TEs expressed in the embryo from those expressed in the endosperm, as well as high reproducibility across biological replicates (Supplemental Figure S1C). In addition, marker genes for embryo or endosperm exhibited their expected expression patterns (Supplemental Figure S1D). Using matched sample pairs from embryo and endosperm collected at 5, 7 and 10 DAP, we drew MA plots to explore the underlying expression differences for TEs. While we detected 105 (5 DAP) and 554 (7 DAP) differentially expressed (DE) TEs during earlier stages of embryo and endosperm development, we identified vastly more DE TEs (*n* = 2,123) in the latest stage (Figure 1, A–C). We subdivided probes into “upregulated in embryo” and “upregulated in endosperm” categories for each tissue (Figure 1, D–F), as TEs may be expressed at low levels in the embryo if they are highly expressed in the endosperm, according to the proposed hypothesis. Although we observed many upregulated TEs in the embryo in our matched sample pairs at 5, 7 and 10 DAP, many upregulated TEs in the endosperm showed a correspondingly low expression level in the embryo (Figure 1, D–F). These results were consistent with the proposed hypothesis that TEs are highly transcribed to silence their counterparts in the embryo genome, although it is also possible that TE expression patterns merely reflect the differences in the developmental program of the embryo and endosperm. In any case, we observed that the expression patterns of TEs were largely distinct between the embryo and the endosperm, as also seen for protein-coding genes (Figure 1, A–C; Supplemental Figure S1D).

**Figure 1.**
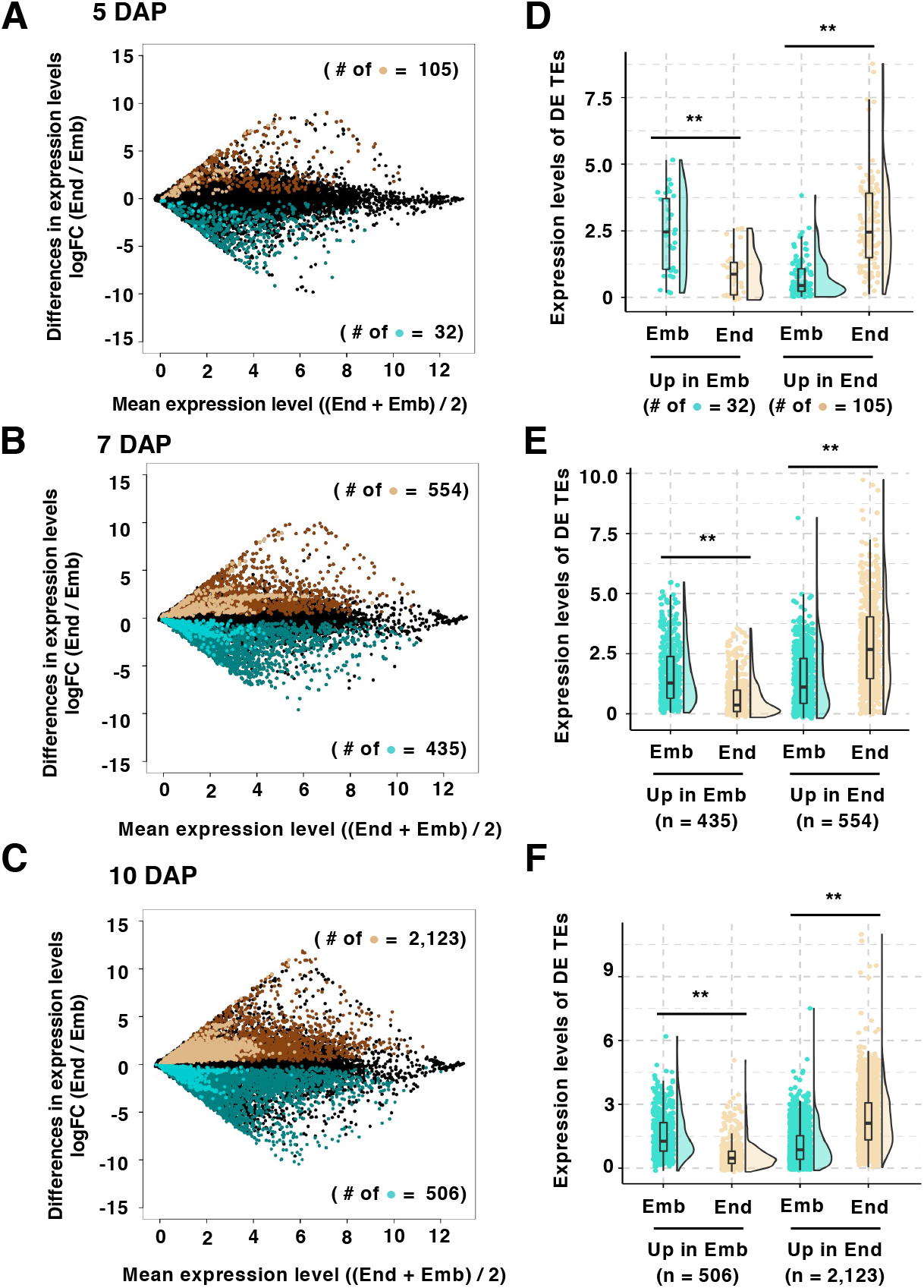
Expression patterns of genes and transposable elements (TEs) in the rice embryo and endosperm. **(A-C)**, MA plots showing the expression levels of genes and TEs at 5 DAP **(A)**, 7 DAP **(B)** and 10 DAP **(C)**. The light and dark brown dots represent differentially expressed (DE) TEs and genes upregulated in the endosperm. The light and dark green dots represent DE TEs and genes upregulated in the embryo. The black dots indicate non-significant probes of TEs and genes (FDR < 0.001). **(D-F)**, Rain cloud plots showing the expression levels of DE TEs. “Up in Emb” and “Up in End” represent negative and positive LogFC dots from MA plots data. The light blue and light brown dots indicate individual expression levels in the embryo and endosperm fractions. **, *P*-value = 0.001; *, *P*-value = 0.05 using a Wilcoxon rank sum test.

### Temporal Expression Patterns of TEs during Rice Endosperm Development

We focused on all samples derived from the endosperm to investigate the temporal expression patterns of the repetitive sequences, including those of TEs, during rice endosperm development from 2 to 10 DAP (Figure 2, A). We determined differentially expressed probes including TEs and repeats during endosperm development by one-way ANOVA and Storey methods (see Materials and Methods) in this data set. We detected dynamic and stage-specific expression patterns when taking a k-means clustering approach, which produced 10 distinct clusters. Clusters A3, A6 and A7 (Figure 2, A) showed an upregulation at 2 and 3 DAP, which may reflect DNA demethylation of the central cell and early endosperm development. However, the majority of repeats appeared to be upregulated at later stages between 7 and 10 DAP, as illustrated byclusters A2, A4, A8, A9 and A10 (Figure 2, A). This result was consistent with the pairwise comparisons of the embryo and endosperm samples at 10 DAP, at which stage more TEs were activated in the endosperm (Figure 1, C and F). Among the DE TEs during endosperm development, the MITE family was prominent in each cluster rather than retrotransposons, DNA transposons, or other repeats (Figure 2, B). We thus extracted the data from MITE probes only for reanalysis (Figure 3, A) and sorted it individual MITE subfamilies (Figure 3, B). As a result, we assessed detectable expression for the MITE subfamilies *Tourist, Castaway, MITE-adh* Type M, *Gaijin/Gaigin* and *Ditto*. These MITE subfamilies were differentially expressed during endosperm development, with broad expression patterns, as all clusters contained representative members of all subfamilies. By contrast, a set of probes corresponding to *Youren* MITEs were predominantly enriched in the M6 cluster (Figure 3, C). *Youren* belongs to the *Tourist*-like elements, with 3-bp (TAA or TTA) target site duplications (Zhang et al., 2004). These elements were characterized by one of the highest signal intensities among all DE TEs in the endosperm at 5 DAP (Figure 3, D) and fairly low signal intensities in the embryo, anthers and leaves (Supplemental Figure S2).

**Figure 2.**
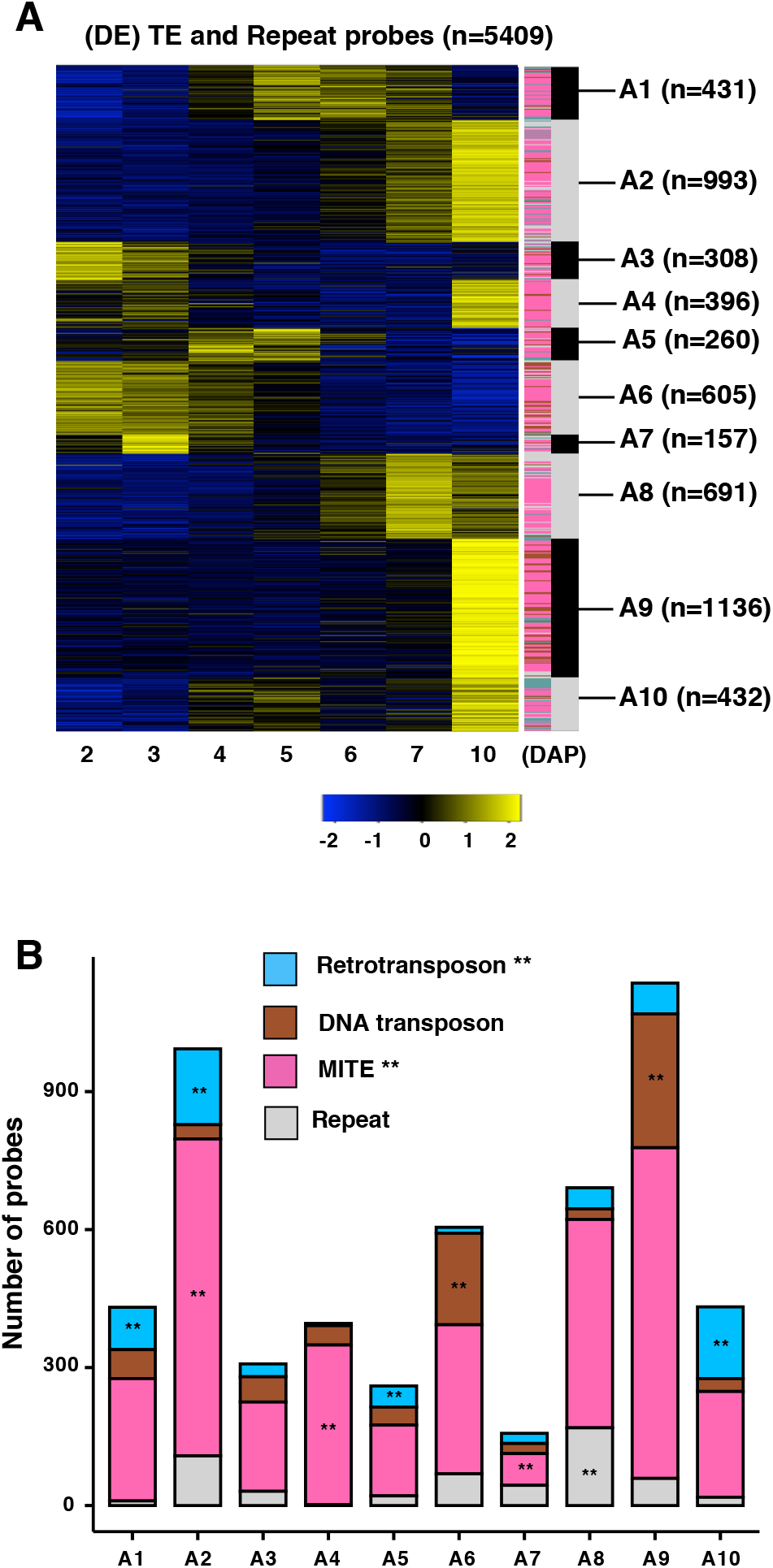
Temporal expression patterns of repetitive sequences. **(A)**, Heatmap representation of k-means clustering results of the transcription patterns of DE TEs and repeats during endosperm development. The colored lines to the right denote representative TE and repeat subclasses; retrotransposons, magenta; DNA transposons, orange; MITEs, deep green; and repeats, gray. The gray and black bars indicate individual clusters. **(B)**, Number of DE probes for the representative subclasses of TEs and repeats in each k-means cluster.

**Figure 3.**
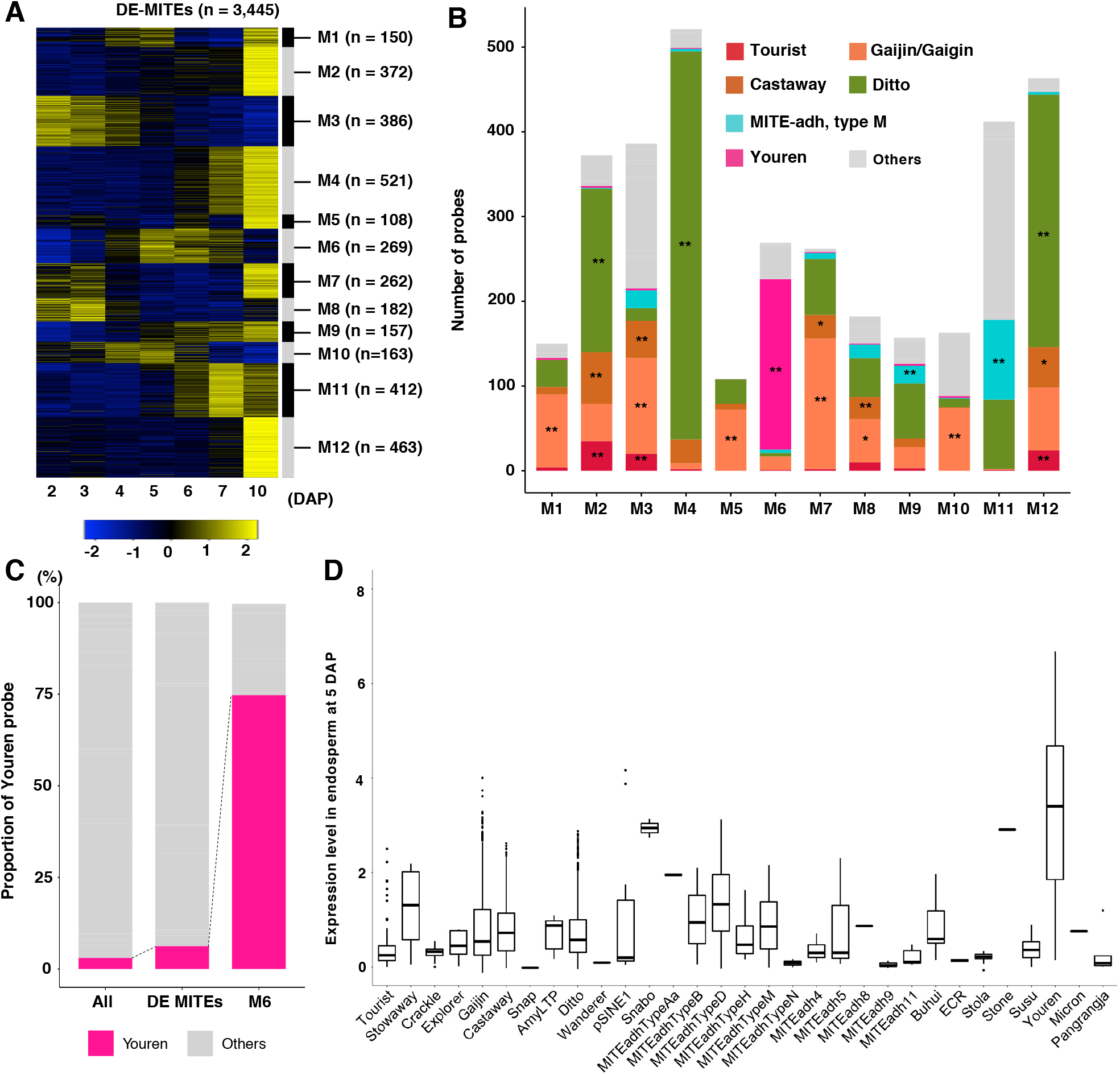
Temporal expression patterns of MITE subfamilies during rice endosperm development. **(A)**, Heatmap representation of row Z-scored expression patterns of DE MITEs during endosperm development. Gray and black bars indicate each cluster from k-means analysis. **(B)**, Number of probes for each representative DE MITE subfamily in individual clusters. **(C)**, Enrichment of the *Youren* subfamily in the M6 cluster among all designed *MITE* and DE *MITE* probes. *Youren* probes, magenta; other MITE subfamilies, gray. **(D)**, Box plots of Log_2_-transformed normalized signal intensity for each MITE subfamily in 5 DAP endosperm. At least one probe for each individual subfamily was differentially expressed in the endosperm from 2 DAP to 10 DAP. The *Youren* subfamily shows one of the highest expression levels.

To assess the possibility of cross-hybridization of highly conserved repetitive transcripts like those derived from TEs, we investigated the relationship between the expression levels and the level of identity between probes (Figure 4, A and B). This possibility was of special concern for *Youren* elements, as they showed overlapping expression patterns but were limited to a single cluster (Figure 3, A and B). We therefore subdivided all *Youren* elements into 11 groups based on their expression level (Figure 4, A) and generated a phylogenetic tree of each probe recognizing each *Youren* element, which revealed no correlation between sequence similarity and signal intensity except for the 10^th^ and 11^th^ weakly expressed groups (Figure 4, B). Therefore, although we do not completely exclude the possibility of cross-hybridization for a small fraction of probes, we conclude that each probe detects transcripts generated from multiple loci in the rice genome.

**Figure 4.**
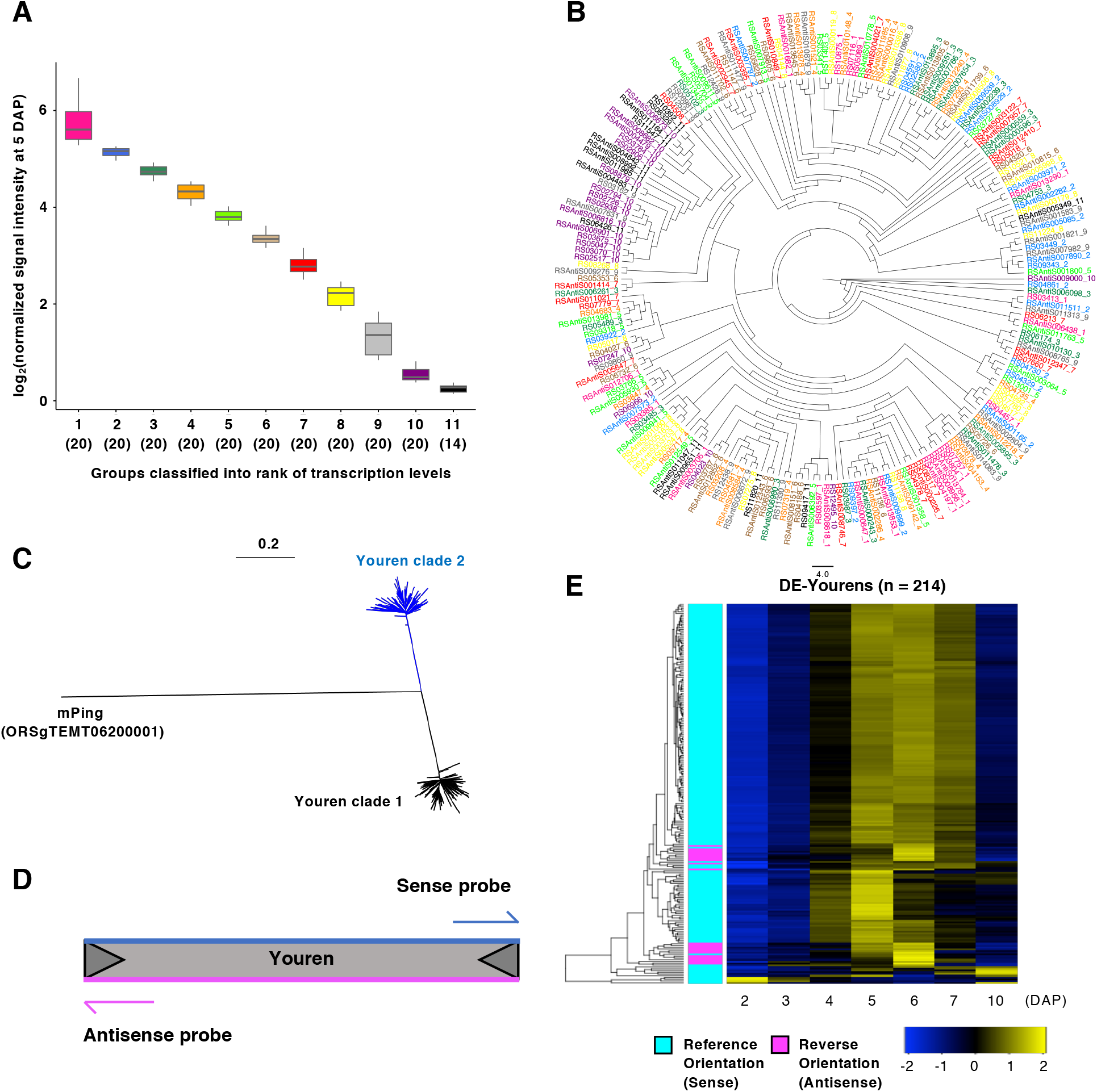
*Youren* TEs are preferentially transcribed in a single direction that matches that of the putative autonomous copy. **(A)**, *Youren* expression levels, arbitrarily divided into 11 groups (214 in total) based on Log_2_-transformed normalized signal intensity in the endosperm at 5 DAP. Numbers in parentheses indicate the number of probes in each group. **(B)**, Sequence similarity of 214 DE *Youren* probes (60-mer), shown as a tree diagram. **(C)**, Phylogenetic tree illustrating the sequence similarity of the entire *Youren* region. Clades 1 and 2 are composed of complementary sequences. **(D)**, Schematic representation of *Youren* TEs and the position of the microarray probes. The Cy-3 labeled complementary RNA is synthesized in reverse direction to the *Youren* transcripts. Therefore, the sense probe detects transcripts in reference orientation (sense strand), whereas the antisense probe detects transcripts in reverse orientation (antisense strand). **(E)**, Heatmap representation of the relationship between expression patterns and transcriptional direction. Color key indicates row Z-score of Log_2_-transformed normalized signal intensity. Cyan, *Youren* in reference orientation; magenta, *Youren* in reverse orientation.

Since MITE sequences are deposited in genome databases without regard for their orientation (Ouyang and Buell, 2004), reference MITE sequences comprise a mixture of forward and reverse orientations (Zhang et al., 2004). Indeed, a phylogenetic tree built from full-length *Youren* sequences in databases formed two distinct clades (Figure 4, C). To determine the direction of *Youren* transcripts, we reannotated each *Youren* probe as reference and reverse orientation, according to the orientation of each *Youren* element deduced from the deposited sequence, taking the direction of transcripts from the putative autonomous copy as a guide. The 60-mer probes printed onto the microarrays were designed to hybridize to the corresponding 5′ and 3′ end sequences of annotated *Youren* elements, making it possible for probes to detect antisense and sense transcripts (Ishiguro et al., 2014) (Figure 4, D). Surprisingly, most *Youren* transcripts appeared to hybridize to the 3′ probes, suggesting that their transcripts are in the reference orientation (Figure 4, D and E). We independently confirmed that *Youren* transcripts are transcribed in the reference strand by RT-PCR (Supplemental Figure S3). Thus, our results suggest the coordinated expression of *Youren* elements originating from different copies scattered over the rice genome. We observed a similar trend, that is, temporal expression patterns were associated with the direction of transcription, for *Tourist, Castaway, MITE-adh* (type M), and *Gaijin* MITE elements, but not for *Ditto* (Supplemental Figure S4).

### Identification of the NF-Y binding motif in *Youren*

We sought to understand the basis for the directional transcription of MITEs. Although we could not distinguish individual insertions of multicopy MITE sequences from the microarray data, we determined locations of all copies of DE TEs and non-DE TEs in the rice genome. The resulting karyoplots indicated that all tested MITE subfamilies map to all chromosomal arm regions, in agreement with a previous report (Mirouze and Vitte, 2014), for both DE MITEs and non-DE MITEs (Supplemental Figure S5). We again focused on highly expressed *Youren* (note that we used 198 different DE *Youren* copies, since eight copies were detected by probes hybridizing to both ends, pointing to these copies as duplications). We also failed to detect any significant skew between MITE subfamilies and DE and non-DE *Youren* elements, for the position of their insertion sites relative to the upstream, coding, intron or downstream region of protein-coding genes (Figure 5, A). The orientations of *Youren* insertions relative to the direction of transcription for their closest gene appeared to be random (Supplemental Figure S6). We thus concluded that these directional transcripts likely arose from internal *Youren* sequences.

**Figure 5.**
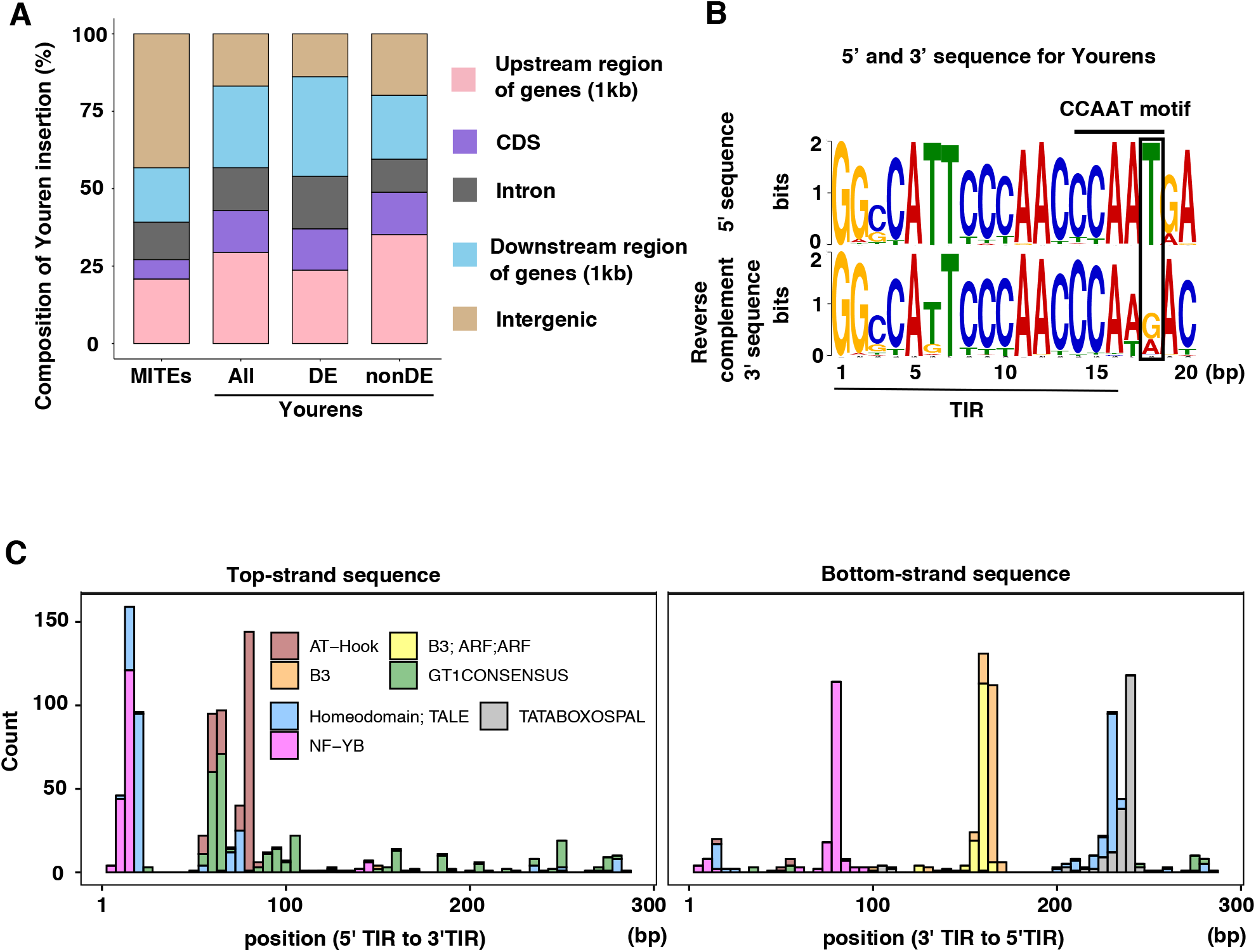
*Youren* insertions in the genome and transcription factor binding motifs. **(A)**, Proportion of insertion sites relative to genes for all MITEs, all *Youren* sequences, as well as DE and non-DE *Youren* sequences. **(B)**, Sequence logos for the 5′ (upper) and 3′ reverse complementary (bottom) sequences that include the TIR and the inner regions, drawn by MEME suite. The horizontal line above the logos indicates the conserved CCAAT motif, known as the NF-Y binding site. The box indicates the last T of the CCAAT motif, which is not conserved in the 3′ sequences of *Youren*. **(C)**, Stacked histogram showing the positions of representative transcription factor binding sites and the top-strand (left) and bottom-strand (right) of DE *Youren. In silico* analysis of transcription factor binding sites of DE *Youren* sequences (*n* = 198) was performed with PlantPAN 3.0 (http://plantpan.itps.ncku.edu.tw).

Since transcript directionality can be attributed to the presence of *cis*-regulatory elements in 5′ sequences, we searched for such elements in all *Youren* copies. A CCAAT motif, which is the typical binding site for NF-Y transcription factors, was present at the boundary between the 5′ TIR and the internal sequence of *Youren*. We also identified a fragment of this motif in the 3′ TIR as a consequence of the inverted repeat, although the internal sequence did not provide the necessary sequence to reconstitute the entire motif (Figure 5, B). We extended our *in silico* analysis to the entire *Youren* sequences using the online tool Plant Promoter Analysis Navigator (PlantPAN 3.0; http://plantpan.itps.ncku.edu.tw), which identified Homeodomain TALE, GT1CONSENSUS, and AT-Hook *cis*-elements (Kyozuka et al., 1994; Boulikas, 1995; Hamant and Pautot, 2010; Xu et al., 2013). Surprisingly, these transcription factor binding sites were preferentially located in the 5′ region of *Youren* for forward orientation, but not for reverse orientation of *Youren* (Figure 5, C) or other tested MITEs. For *Youren* elements with detectable transcripts by RT-PCR, the AT-Hook binding sites located within the forward primer used for PCR, suggesting that the element overlaps with the transcribed *Youren* region (Supplemental Figure S3). The NF-Y binding motif is a good candidate to explain the directional transcription of *Youren*, as several genes encoding the trimeric NF-Y complex, *NF-YB1, NF-YB7*, and *NF-YB9*, are predominantly expressed in the aleurone and starchy endosperm and control seed development in rice (Bai et al., 2016; Niu et al., 2020; Niu et al., 2021). In addition, the temporal expression patterns of *NF-YB1, NF-YB7*, and *NF-YB9* are similar to those of *Youren* elements. In conclusion, we identified coordinated expression patterns of *Youren*s and the potential underlying *cis*-elements.

## DISCUSSION

Hypomethylation of the genome is observed in the endosperm of several plant species, which prompted us to investigate the expression patterns of rice TEs during endosperm development and the epigenetic control mechanisms of TE expression in relation to protein-coding gene expression and genome integrity. TEs can influence the expression of their neighboring gene, either by preventing transcription or by promoting transcription via providing an auspicious chromatin environment for active transcription (Feschotte, 2008; Qiu and Kohler, 2020). In addition, some transcripts derived from TEs exhibit a sponge-like activity against small RNAs and miRNAs through hybridization and sequestration away from their targets (Cho and Paszkowski, 2017). In other cases, transcripts from MITEs provide the stem structure required to form the “stem-loop” structure of pre-miRNAs. Therefore, temporal analysis of TE expression originating from hypomethylated endosperm is an ideal resource to study the roles of TEs.

Since genome-wide demethylation has been observed in the central cell and endosperm cells, a simple prediction is that TE activity would be high in the endosperm at all developmental stages (Ibarra et al., 2012; Park et al., 2016). However, our temporal analysis revealed a very different picture, as the expression of TEs was not static and high but exhibited dynamic changes, with some TEs expressed during earlier stages and others expressed later. We observed a similar trend in the embryo, although we limited our investigation to three time points, representing the coleoptile, the completion of organogenesis and the maturation stages of rice embryogenesis. Over the course of these time points, the embryo develops into a mature structure that consists of several tissues: the root and shoot primordium, leaves and the scutellum. The expression of rice TEs was reported to be largely different in distinct plant tissues (Cho and Paszkowski, 2017); our results therefore likely merely reflect the tissue- and organ-specific transcriptional programs in the embryo. Similarly, the rice endosperm develops into several domains, such as the peripheral region and the region adjacent to the embryo, as well as the aleurone layer and the starchy endosperm, and the expression profile of TEs may differ in each compartment. Alternatively, the transcriptional environment of TEs may rely on temporal programs and epigenetic changes of TE sequences. Demethylation of the maternal genome has been observed in several plant species, resulting in imprinted gene expression (Hsieh et al., 2009; Rodrigues et al., 2013; Batista and Kohler, 2020). *Youren* has the potential to contribute to this type of epigenetic modulation of gene expression (Yuan et al., 2017). In addition to the demethylation of the maternal copy of the endosperm genome, DNA methylation status changes during seed and endosperm development in both Arabidopsis and rice (Xing et al., 2015; Kawakatsu et al., 2017). To better understand the roles of TEs and demethylation in gene expression during endosperm development, it is necessary to conduct higher-resolution expression analyses at the cell type level, combined with multiple omics analyses to uncover the epigenetic states underlying endosperm development.

Although the transcription of long and almost intact TEs like LTR retrotransposons is directionally controlled, the transcription of remnant TEs is thought to be influenced by the expression of neighboring genes and may even form chimeric transcripts bringing together gene sequences and TEs. MITEs are remnant TEs derived from longer autonomous copies and thus no longer encode the sequences required for their transposition. Nevertheless, transcripts derived from MITE loci, with the exception of *Ditto*, are directionally controlled. Our results suggest that these elements might have preserved the promoter activity required for autonomous transcription. Among the tested MITEs, *Youren* was unique, as the expression of most elements was relatively high, similar and directionally controlled. We hypothesize that these patterns are caused by *cis*-elements that reside in the 5′ internal region of *Youren*, since the sequences outside of TE insertion sites are not conserved and cannot explain their coordinated expression. The approach we followed in this study provides the framework to test our proposed hypothesis. TEs provide *cis*-elements that can therefore rewire transcriptional networks (Feschotte, 2008). Although direct binding and transcriptional control will need to be tested, the trimeric NF-Y complex is good candidate. The NF-YB and NF-YC subunits have histone-fold domains to bind to DNA in a non-sequence-specific manner, while NF-YA recognizes the CCAAT motif (Nardini et al., 2013). Although pioneer transcription factors are not known in the endosperm, Arabidopsis LEAFY COTYLEDON1 (LEC1), an NF-YB subunit, is involved in embryogenesis (Tao et al., 2017). We identified 198 *Youren* copies actively transcribed during the mid-stage of endosperm development, the timing of which is consistent with the onset of transcription for genes encoding starch synthases and storage proteins (Tonosaki et al., 2021). These copies of *Youren* are inserted in 700 distinct locations in the rice genome close to genes and provide potential *cis*-elements for trimeric NF-Y binding. It will be worth investigating whether and how much these elements can rewire transcriptional networks during endosperm in the context of the evolutionary history of rice.

## MATERIALS AND METHODS

### Plant Materials and Growth Conditions

Rice plants (*Oryza sativa* cv. Nipponbare) were grown in a Biotron chamber (NC-350HC; Nihon Ikakikai) according to a previously described protocol (Ohnishi et al., 2011) adjusted for short-day photoperiod and temperature cycles (11-h light at 30°C/13-h dark at 25°C).

### RNA Extraction from Rice Endosperm

Rice endosperm was harvested in 24-h intervals from developing seeds from 2 to 7 d after pollination (DAP) and at 10 DAP (Supplemental Figure S1A). For the isolation of liquid coenocytic endosperm (2 DAP) and just after cellularization (3 DAP), a cut was made on the surface of the seed coat using a scalpel under a binocular microscope, as described previously (Tonosaki et al., 2021). The liquid endosperm was collected from the cut by pressing on the seed using a pipette and added to 50 µL of the extraction buffer from the PicoPure RNA isolation kit (Applied Biosystems). Between 70 and 100 seeds at 2 DAP and 50 seeds at 3 DAP were used for isolation of liquid endosperm. After 4 DAP, endosperm tissues were directly dissected from the ovaries under a binocular microscope, followed by homogenization in RNA isolation cocktail, consisting of 350 µL of RLC buffer (Plant RNeasy kit; Qiagen), 100 µL of Fruit-mate (Takara) and 4.5 µL of β-mercaptoethanol (Wako) (Lee et al., 2016) using micro pestles and 1.5-mL plastic tubes. Between 10 and 30 ovaries were used for isolation of endosperm from 4 to 7 DAP. Total RNA was extracted with the RNeasy Plant Mini Kit according to the manufacturer’s instructions (Qiagen). At 10 DAP, dissected endosperm tissue from three ovaries was directly transferred to 1.5-mL plastic tubes and immediately frozen in liquid nitrogen. The frozen endosperm was homogenized with a multi-beads shocker (MB755U, Yasui Kikai) at 1,500 rpm for 2 s. The resulting homogenized endosperm was resuspended in 1.3 mL of Fruit-mate. The mixture was subdivided into two 650-µL aliquots and centrifuged at 12,000*g* for 5 min at 4°C. To 250 µL of each supernatant, 400 µL of RLT buffer from the RNeasy Plant Mini Kit (Qiagen) was added together with 4 µL of β-mercaptoethanol (Wako). For DNase treatment, RNase-Free DNase Set (Qiagen) was used according to the manufacturer’s instructions.

### RNA Extraction from Rice Embryos

Rice embryos were dissected from developing ovaries at 5, 7 and 10 DAP (Supplemental Figure S1B) under a binocular microscope. Since early-stage embryos are small and difficult to dissect under a binocular microscope, two protocols were tested to isolate RNA from embryos at 5 DAP: the RNeasy Plant Mini Kit and the PicoPure RNA isolation kit (Thermo Fischer Science). For the RNeasy Plant Mini Kit, 15 embryos were collected from developing ovaries and were immediately frozen in liquid nitrogen. The frozen embryos were homogenized with a multi-beads shocker at 1,500 rpm for 2 s; the subsequent steps followed the protocol described above. For the PicoPure RNA isolation kit (Thermo Fischer Science), three embryos at 5 DAP were homogenized in 10 µL of extraction buffer from the PicoPure RNA isolation kit using a micro pestle and a 1.5-mL microcentrifuge tube. The homogenate was incubated at 42°C for 30 min. Total RNA was extracted according to the manufacturer’s instructions. At 7 and 10 DAP, three embryos were collected for extraction of total RNA using the RNeasy Plant Mini Kit. The quality of RNA samples was assessed on an Agilent 2100 Bioanalyzer (Agilent Technologies)

### Microarray Analysis

The 44k Agilent microarray slides used in this study were prepared according to a previously published method (Ishiguro et al., 2014). The labeling reactions were performed using a Quick Amp Labeling kit (Agilent). The Cy3-labeled cRNAs were fragmented and hybridized to the microarray slides at 65°C for 17 h. One-color spike mix was added to total RNA samples before reactions and used to monitor the efficiency of the labeling reactions and hybridizations. Hybridization and washing conditions of the hybridized slides followed the Agilent protocol. The slides were scanned on an Agilent G2565BA microarray scanner (Agilent). The background correction of raw Cy3 signals was conducted using the Agilent Feature Extraction software (version 9.5.1; Agilent). All microarray analyses were performed with three independent biological replicates of embryo and endosperm tissues.

### Microarray data analysis

Data analysis was performed in R (version 3.5.2). To determine the difference in TE expression between 5, 7 and 10 DAP embryos and endosperm, the raw signal data were normalized by the Quantile method in the R package *limma* (version 3.38.3)(Ritchie et al., 2015). The individual log_2_-transformed normalized signal densities that were the mean of signals of negative-control probes were used as expression levels. Differentially expressed probes between the embryo and endosperm at 5, 7 and 10 DAP were identified with the row *t*-test in the R package *genefilter* (version 1.64.0) (Gentleman et al., 2021) and with the Storey Tibshirani method in the R package *qvalue* (version 2.14.1) (Storey et al., 2020). The false discovery rate (FDR) was set to 0.001. To analyze the differences in TE expression during endosperm development, raw data from 2 to 7 DAP and 10 DAP samples were normalized by the Quantile method. Differentially expressed probes during endosperm development were identified by one-way ANOVA and Storey methods with a FDR of 0.001. The number of k-means clusters of differentially expressed TEs, retrotransposons, DNA transposons and MITEs was determined based on the k-gap score calculated by clusGap in the R package *cluster* (version 2.1.0) (Yao, 2013; Maechler et al., 2021). For comparison of the levels of *Youren* transcripts across rice tissues, the signal densities from leaves and anthers were retrieved from the microarray data series GSE49561 from the Gene Expression Omnibus database at NCBI. The alignment of DE *Youren* probes was performed in ClustalW and the resulting phylogenetic tree was drawn with FigTree. The genomic distribution of MITEs was plotted with the R packages *karyoploteR* (version 1.16.0) (Gel and Serra, 2017) and *BSgenome*.*Osativa*.*MSU*.*MSU7* (version 0.99.1)(Yao, 2013).

### RT-PCR analysis of Youren

First-strand cDNA was synthesized with the PrimeScript First Strand cDNA Synthesis Kit (Takara) with oligo-dT primers and total RNA isolated from 5 DAP endosperm. Oligo-dT was chosen here to match the conditions used for microarray labeling. First-strand cDNAs were used as template for PCR with the forward primer F1 (5′-CACCGGCAAGTGAATAAATGAGGAAA*<u>T</u>*-3′) and the reverse primer R (5′-TCCCAACCCAWRACACTAGACAT-3′), which were designed to anneal to *Youren* copy with the highest transcript level in 5 DAP endosperm. PCR reactions were performed with SeqAmp DNA Polymerase (Takara), a proofreading polymerase. The 3’ end sequence of the F1 primer was set to one of the single nucleotide polymorphism (SNP) sites of the *Youren* family to specifically amplify the most highly expressed copy (three copies in the rice genome). For the other *Youren* copies, the F2 primer (5′-CACCGGCAAGTGAATAAATGAGGAAA*<u>G</u>*-3′) was used instead, with PCR reactions using FastStart Taq DNA Polymerase (Sigma-Aldrich), which does not have proofreading activity. Therefore, the SNP site cannot not be corrected during the amplification, allowing the amplification of other copies expressed at lower levels. *ACTIN* was used as a control for RT-PCR (Ishikawa et al., 2011). *Youren* RT-PCR products were cloned into the pENTR™/D-TOPO™ vector (Takara) and sequenced by Sanger sequencing (Applied Biosystems 3500).

## Supplemental Data

The following supplemental materials are available.

**Supplemental Figure S1**. Expression analysis of genes and TEs during endosperm development.

**Supplemental Figure S2**. Expression levels of *Youren* elements are high in the endosperm.

**Supplemental Figure S3**. RT-PCR analysis of *Youren* transcripts.

**Supplemental Figure S4**. Temporal expression patterns and transcriptional direction of *MITE* subfamilies.

**Supplemental Figure S5**. Distribution of *Youren, Tourist, Castaway, MITE*-adh (type M), *Gaijin* and *Ditto* copies corresponding to the microarray probes.

**Supplemental Figure S6**. Proportion of *Youren* orientation relative to the nearest transcriptional unit.

## ACKNOWLEDGMENTS

This work was partly supported by a Grand-in-Aid for Scientific Research on Innovative Area (16H06464, 16H06471 and 16H21727 to T.Ki., 19H04873 to T. Ka.) and a JSPS Research Fellowship (16J02580 to K.T.) from the Ministry of Education, Culture, Sports, Science and Technology (MEXT) of Japan.

## AUTHOR CONTRIBUTIONS

K.T., A.O. and T.K. conceived research plans; T.K. and L.C. supervised the experiments; K.T., H.N., M.K. and T.K. performed the experiments; K.T analyzed the data; H.F., K.N., Y.S. and M.E. provided research materials; K.T. and T.K. wrote the paper.

